# Using TrackMate to Analyze *Drosophila* Larval and Adult Locomotion

**DOI:** 10.1101/2021.09.28.462241

**Authors:** Alisa A. Omelchenko, Ainul Huda, Allison N. Castaneda, Thomas J. Vaden, Lina Ni

## Abstract

*Drosophila* adult and larvae exhibit sophisticated behaviors that are widely used in development, synaptic transmission, sensory physiology, and learning and memory research. Many of these behaviors depend on locomotion, the ability of an animal to move. However, the statistical analysis of locomotion is not trivial. Here we use an open-source Fiji plugin TrackMate to track the locomotion of *Drosophila* adults and larvae. We build optimal experimental setups to rapidly process recordings by Fiji and analyze by TrackMate. We also provide tips for analyzing non-optimal recordings. TrackMate extracts the X and Y positions of an animal on each frame of an image sequence or a video. This information allows for generating moving trajectories, calculating moving distances, and determining preference indices in two-choice assays. Notably, this free-cost analysis method does not require programming skills.

**Summary statement:** This study uses an open-source Fiji plugin TrackMate to computationally analyze *Drosophila* adult and larval behavioral assays, which does not require programming skills.

## Introduction

*Drosophila melanogaster* exhibit sophisticated behaviors that are widely used in studies of development, synaptic transmission, sensory physiology, and learning and memory. Many of these behaviors depend on locomotion, the ability of an animal to move. The locomotion analysis of larvae and adult flies is essential to gather insights into how modifications of genetic components affect animal behaviors and responses to stimuli. The examination of their movement thereby has become an integral part of such studies leading to the development of tracking systems to provide a quantitative description of their behaviors (Bellen et al., 2010).

Several tracking systems have been developed to track the locomotion of larvae and adult flies and generate trajectories (Werkhoven et al., 2019, Branson et al., 2009, Valente et al., 2007, Straw and Dickinson, 2009, Colomb et al., 2012). However, many methods require programming skills and/or commercial software to set up and run the tracking systems, which often become obstacles for researchers to adopt these methods. Other methods require specific experimental setups to collect data and are challenging to analyze non-optimal recordings. Therefore, it is not trivial to statistically analyze the locomotion of larvae and adult flies. There is a need to develop an open-source approach that does not require programming skills but can computationally track the movement of larvae and adult flies and generate trajectories. Ideally, this method can analyze non-optimal recordings.

In this study, we use an open-source Fiji plugin TrackMate to achieve this goal. We build optimal experimental setups to rapidly process and analyze behavioral recordings. We also provide examples of analyzing non-optimal recordings. TrackMate extracts the X and Y positions of an animal on each frame. This information allows for generating moving trajectories, calculating moving distances, and determining preference indices in two-choice assays. Notably, this free-cost analysis method does not require programming skills. This method is validated by analyzing the free motion and two-choice thermotactic behavioral discrepancies of wild type and thermoreceptor mutants in larvae and adult flies.

## Results

### Using TrackMate to analyze the locomotion of adult flies

We first used TrackMate to analyze the locomotion of adult flies. Since TrackMate recognizes particles (regions of interest (ROIs)) based on their intensities, background noise signals can be recognized and mistakenly tracked as ROIs. We suggested maximally diminishing the background noise signals by performing the assay on a piece of white paper (**Fig. 1A**). In our setup, the fly was covered by a transparent cover that only allowed the fly to walk, not fly. A Styrofoam box was placed to cover the experimental region to create a featureless environment with dim ambient light of < 10 lux. A GoPro camera was installed on the ceiling in the Styrofoam box to take time-lapse pictures (1 picture per second) for 2 minutes. These pictures were then imported into Fiji as an image sequence and preprocessed. Next, TrackMate was run to track the fly movement, extract its X and Y positions, generate the trajectory (**Fig. 1C**), and the moving distance was calculated (**Fig. 1D**). We found that the warm receptor mutant, *Gr28b*^*MB*^, moved significantly more than *wild-type* (*wt*) flies (**Fig. 1C,D; Movie 1**).

**Fig 1.**
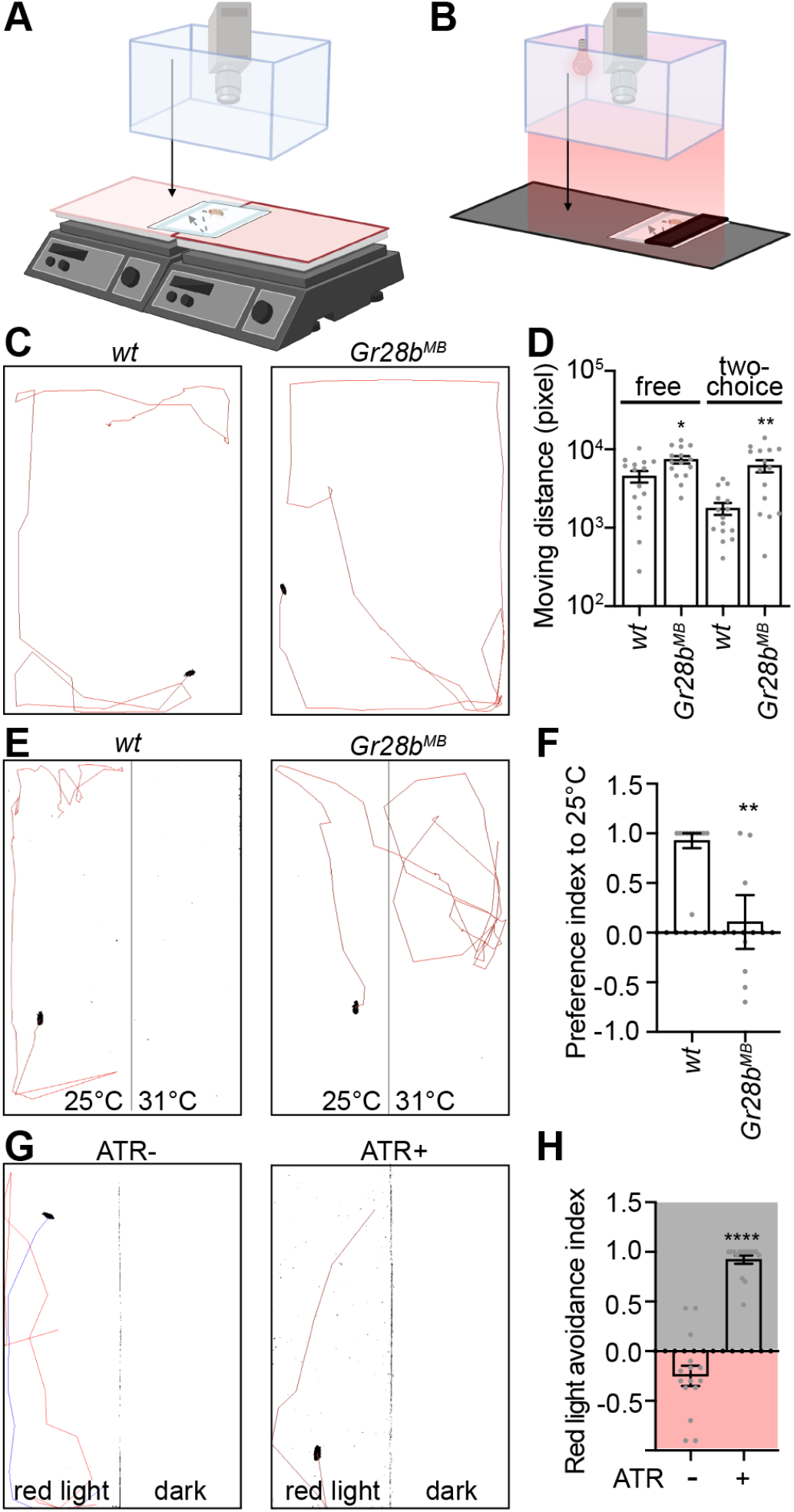
Use TrackMate to analyze the adult behavior. (A,B) Setups for the single-fly two-choice thermotactic (A) and optogenetic assay (B). The setup for the free motion assay is similar to the two-choice assay but is performed on a single plate with a unique temperature (25°C). (C) *wt* and *Gr28b*^*MB*^ trajectories in the free motion assay. (D) Moving distances of indicated genotypes and conditions. n = 15; data represent mean ± SEM; * *p* < 0.05, ** *p* < 0.01; Welch’s test for the free motion assay and Mann-Whitney test for the two-choice assay. (E) *wt* and *Gr28b*^*MB*^ trajectories in the two-choice assay. (F) Preference indices of indicated genotypes. n = 7 - 11; data represent mean ± SEM; ** *p* < 0.01; Mann-Whitney test. (G) Trajectories of *HC-Gal4;UAS-CsChrimson* (*HC*>*CsChrimson*) flies with or without dietary retinal (ATR) in the optogenetic assay. Of note, two trajectories (blue and red) are shown in the left panel, suggesting the fly enters the red-light zone twice. (H) Avoidance indices of *HC*>*CsChrimson* flies with or without dietary retinal (ATR). n = 15; data represent mean ± SEM; **** *p* < 0.0001; Mann-Whitney test.

TrackMate was also used to analyze the two-choice assay. In the two-choice assay, a line was drawn to separate different temperatures (**Fig. 1E**). According to the X position of the line and the X position of the fly on each frame, the time the fly spent in each temperature zone was calculated and the preference index was determined. While *wt* flies avoided 31°C and preferred 25°C, *Gr28b*^*MB*^ did not show preference between 25°C and 31°C (**Fig. 1F**). This result is consistent with previous reports (Ni et al., 2013, Simões et al., 2021, Budelli et al., 2019). Moreover, *Gr28b*^*MB*^ also moved more than *wild-type* (*wt*) flies in this condition (**Fig. 1D,E; Movie 1**).

An optogenetic assay was also analyzed by TrackMate. The optogenetic tool used in this study was the red light-shifted channelrhodopsin CsChrimson. When bound to all-trans retinal (ATR), CsChrimson is activated by the red light to depolarize cells (Klapoetke et al., 2014). To create the dark and red-light environments, a red-light source was placed on the ceiling in the Styrofoam box and half of the transparent cover was covered by black tape (**Fig. 1B**). Of note, the red light must not be directly above the cover (See the section of Using TrackMate to analyze nonoptimal recordings and **Fig. 3**). We suggested installing the GoPro camera in the middle of the Styrofoam box, the light source on its left, and the experimental region at the right of the Styrofoam box (**Fig. 1B**). The recording procedure and analysis method were similar to the two-choice assay. Heating Cells (HCs) in aristae drive warm avoidance (Gallio et al., 2011, Budelli et al., 2019, Ni et al., 2013). Flies expressing CsChrimson in HCs avoided red light with dietary ATR (**Fig. 1G,H; Movie 1**). Without ATR, HCs weren’t activated and did not guide flies to avoid the red light. This was validated by the observation that flies often traveled from the dark zone to the red-light zone and thus the red-light zone had multiple tracks (**Fig. 1G; Movie 1**).

### Using TrackMate to analyze the larval locomotion

Next, we used TrackMate to track the larval movement and analyzed the larval two-choice thermotactic assay. The larva was allowed to move on 3% agar gel. To diminish the background noise signals and increase contrast, a sheet of matte black poster was placed under the gel and ambient light was dimed to under 10 lux. A GoPro camera was placed above the gel to record the larval movement for 2 min. For the two-choice assay, the larva was released in the middle of two temperatures (**Fig. 2A**). The 2-min video was imported into Fiji and preprocessed. For the two-choice assay, a line was drawn to separate different temperatures. Of note, this line must not pass the moving path of the larva (**Fig. 2E**). TrackMate was then used to track the larval movement and generate its trajectory (**Fig. 2C,E**). TrackMate also extracted X and Y positions of the larva on each frame to calculate its moving distance and preference index (**Fig. 2D,F**).

**Fig 2.**
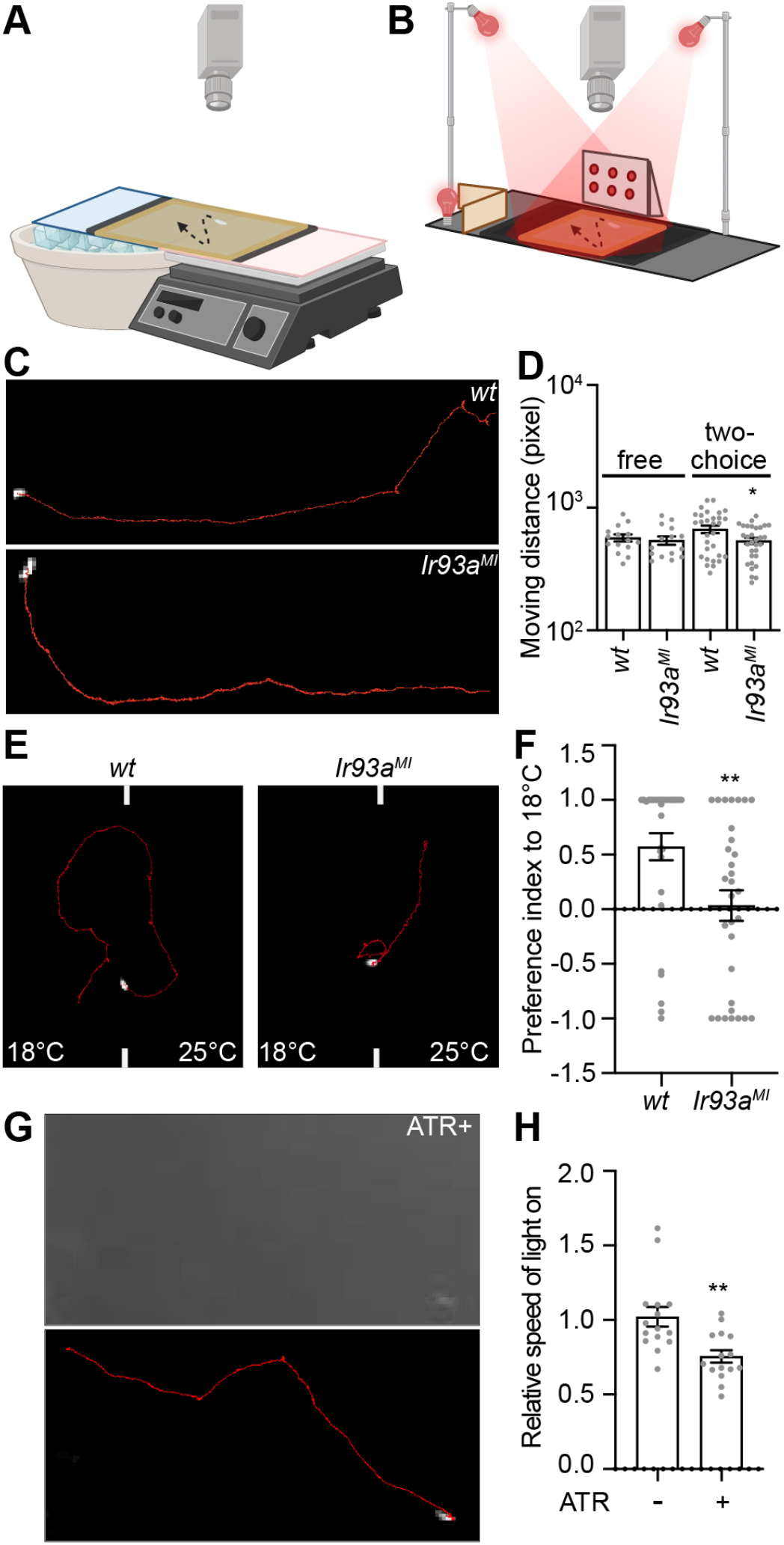
Use TrackMate to analyze the larval behavior. (A,B) Setups for the single-larva two-choice thermotactic (A) and optogenetic assay (B). The setup for the free motion assay is similar to the optogenetic assay but is performed in the room light. (C) *wt* and *Ir93a*^*MI*^ trajectories in the free motion assay. (D) Moving distances of indicated genotypes and conditions. n = 15 - 30; data represent mean ± SEM; * *p* < 0.05; Welch’s test for the free motion assay and Mann-Whitney test for the two-choice assay. (E) *wt* and *Ir93a*^*MI*^ trajectories in the two-choice assay. (F) Preference indices of indicated genotypes. n = 30; data represent mean ± SEM; ** *p* < 0.01; Mann-Whitney test. (G) The locomotion of an *Ir21a-Gal4;UAS-CsChrimson* (*Ir21a*>*CsChrimson*) larva with ATR in the optogenetic assay. Upper: the video is gray scaled and cropped by Fiji. Lower: the recording is preprocessed by Fiji and analyzed by TrackMate. (H) The relative speed of *Ir21a*>*CsChrimson* larvae with or without ATR. Relative speed is defined as the moving speed during red light on divided by the moving speed during light off. N = 15; data represent mean ± SEM; ** *p* < 0.01; Mann-Whitney test.

IR93a is a subunit of the cool receptor in dorsal organ cool cells (DOCCs) and guides larvae to avoid cool temperatures (Knecht et al., 2016). In a unique temperature environment, moving distances of *wt* and *Ir93a*^*MI*^ larvae were similar (**Fig. 2C,D; Movie 1**). However, *Ir93a*^*MI*^ larvae moved significantly less than *wt* larvae in a two-temperature environment (**Fig. 2D,E; Movie 1**). Regarding preference indices, *wt* larvae preferred 18°C (**Fig. 2E,F; Movie 1**). This is consistent with previous reports (Kwon et al., 2008, Kwon et al., 2010, Shen et al., 2011). The *Ir93a*^*MI*^ larvae had no preference between 18°C and 25°C, suggesting IR93a is required for choosing the optimal temperature within this temperature range (**Fig. 2E,F; Movie 1**).

The larval optogenetic assay must be performed under an infrared condition while avoiding light glares. The red-light intensity should be even across the region where the larva travels (**Fig. 2B**). The recording procedure and analysis method were similar to the free motion assay. Larvae expressing *CsChrimson* in DOCCs showed aversive behaviors under red light with dietary ATR, such as the pause of run, which in turn led to the decrease of the run speed (**Fig. 2G,H**) (Tyrrell et al., 2021). These aversive behaviors reflected the cool avoidance driven by DOCCs and were not observed in the group without ATR (**Fig. 2G,H**).

### Using TrackMate to analyze non-optimal recordings

If behavioral recordings are available and computational analysis is required, TrackMate is an option. The analysis process may take longer if the recordings contain background noise signals. **Fig. 3A** showed a setup for optogenetics previously used in the lab (Tyrrell et al., 2021). The light source was placed under an agar plate so the light source and light glares were recorded (**Fig. 3A**). After being gray scaled, the larva was detected but a significant amount of background noise signals were also shown (**Fig. 3B**; the new setup had a cleaner background shown in **Fig. 2G**). We suggested cropping the video and using the smallest possible region for analysis. Fiji parameters, such as background, brightness, and contrast, must be adjusted to diminish background noise signals. In most cases, not all background noise signals could be avoided (circle in **Fig. 3B**). TrackMate parameters, including LoG detector, filters on spots, simple LAP tracker, and filters on tracks, must also be optimized to detect the larva in most frames (>99%) and maximally decrease noise signals. Finally, the All Spots statistics.csv file must be carefully checked to ensure all ROIs related the larva, not noise signals. The colorful track suggested the larva was not detected or counted more than once on some frames so that multiple tracks were generated (**Fig. 3B**). Although having taken longer to process and analyze, these non-optimal recordings found similar results that DOCC expression of CsChrimson drove aversive behaviors (**Fig. 3C**).

**Fig 3.**
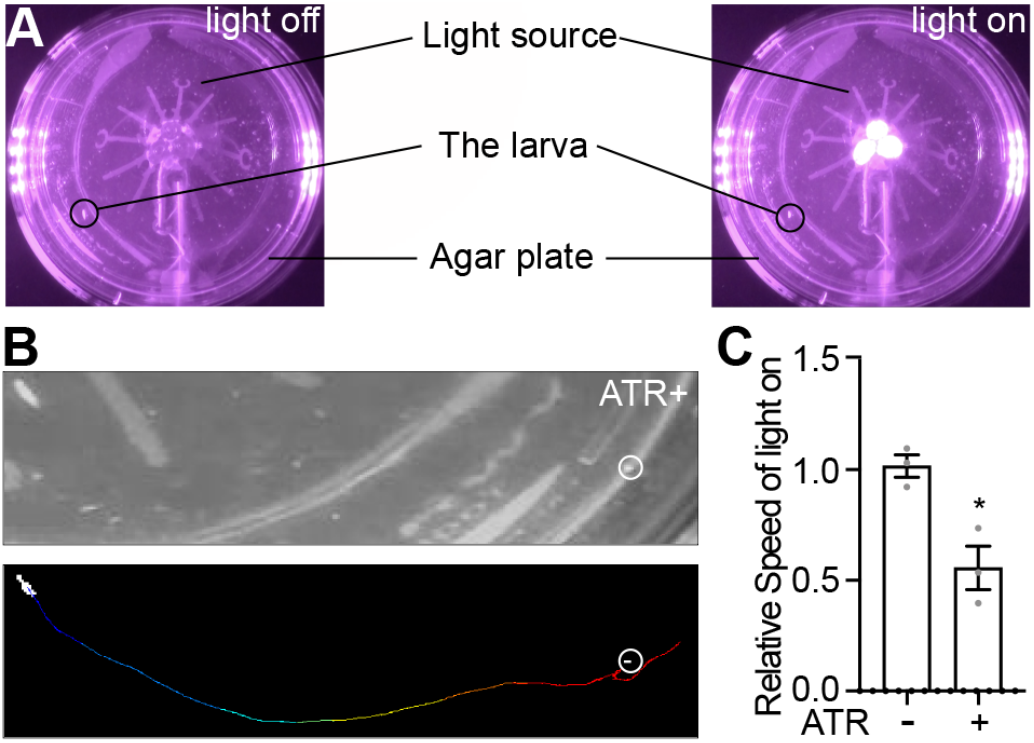
Use TrackMate to analyze non-optimal recordings. (A) A previous setup of the single-larva optogenetic assay, in which light glares and other background noise signals are detected. (B) The locomotion of an *Ir21a*>*CsChrimson* larva with dietary retinal (ATR). Upper: the video is gray scaled and cropped by Fiji. Lower: the recording is preprocessed by Fiji and analyzed by TrackMate. The trajectory is shown in rainbow colors. (C) The relative speed of *Ir21a*>*CsChrimson* larvae with or without ATR. n = 3; data represent mean ± SEM; * *p* < 0.05; Welch’s test.

## Discussion

In this study, we apply an open-source Fiji plugin TrackMate to track the locomotion of adult flies and larvae in image sequences and videos and generate trajectories. Since TrackMate extracts the X and Y positions of an animal on each frame, it is used to analyze preference indices in two-choice assays, in which the separating line can be defined as X or Y positions. In addition, TrackMate can be used to analyze non-optimal recordings with strong background noise signals.

Clean background facilitates the analysis. TrackMate detects ROIs – flies or larvae – based on their intensities. It cannot distinguish ROIs from background noise signals if they have similar sizes and/or intensities. Thereby, noise signals cause aberrant trajectories and require researchers to adjust TrackMate parameters to avoid these signals or check the All Spots statistics.csv file to delete information for noise signals in the file. We provide easy ways to get clean backgrounds – a piece of white paper and matte black poster sheet can significantly decrease background noise signals in adult and larval behavioral assays, respectively.

TrackMate extracts the X and Y positions of the ROI on each frame. According to this information, the moving distance is determined, calculated in pixels. If moving distances from two experiments need comparing, it is essential to convert the pixel distance to the actual distance. Moreover, the camera position and image size must be kept constant from trial to trial to avoid the conversion. To keep the analysis simple and accurate, we recommend using the same setup to perform all experiments that need comparing.

TrackMate can analyze data recorded in image sequences and videos. In this study, we use time-lapse images to record adult behaviors and videos to record larval behaviors. Thereby, analysis approaches for both recording methods have been shown. In videos, every second contains 24-30 frames; their high temporal resolutions can resolve more behavioral details. When calculating moving distances and preference indices, high temporal resolutions are not necessary. In this case, time-lapse images become a better choice because they contain less data and take a shorter time to analyze. However, the time-lapse image resolution of the GoPro cannot distinguish the larva from the background and videos are used for larval assays instead. When performing the larval optogenetic assay, we find it is easier to track the larva under infrared conditions. Of note, regular cameras don’t work under infrared conditions – the internal infrared filter needs to be removed and replaced by an 830 nm long-pass filter.

Based on trajectories, researchers can observe whether an animal runs or turns. But TrackMate may not be used to analyze more complex behaviors, such as larval head sweeping. We set a relatively large blob diameter to make TrackMate recognize the adult fly or larva as a single ROI. If using a relatively small blob diameter, TrackMate often detects the animal as multiple ROIs. But the localization of each ROI cannot be predicted; in other words, TrackMate cannot be used to recognize different parts of an animal, such as the head or tail. Thereby, it is challenging to use TrackMate to analyze more complex behaviors.

## Materials and methods

### *Drosophila* strains

*CS* and *WCS* were used as wild-type controls for larval and adult behavioral assays, respectively. The following flies were previously described: *UAS-CsChrimson* (Klapoetke et al., 2014), *Ir93a*^*MI*^ (Knecht et al., 2016), *Ir93a-Gal4* (Sanchez-Alcaniz et al., 2018), *Gr28b*^*MB*^ (Ni et al., 2013), *HC-Gal4* (Gallio et al., 2011).

### Adult behavioral assays

Flies were raised at 25°C under 12-hour light/12-hour dark cycles and were 3-7 days old when tested. For the two-choice thermotactic assay, two steel plates on different hot plates were aligned so that the steel plate boundaries were brought together. Hotplate temperatures were adjusted allowing the surface of the steel plates to be 25± 1°C and 31± 1°C, respectively. For the free movement behavioral assay, a steel plate was placed on a hot plate and its surface temperature was adjusted to 25± 1°C. The plates were sprayed with dH_2_O and a plastic sheet protector was placed on top. Excess moisture was removed by Kimwipe. A white paper was placed on top of the plastic sheet protector to reduce background noise signals. A clear plastic cover was covered with SigmaCote to prevent the fly from walking on the plastic cover and was placed on top of the white sheet so that it was divided evenly on the steel plate boundary creating the experimental chamber. The temperature was monitored before each trial using a surface temperature probe (80PK-3A, Fluke) and thermometer (Fisherbrand Traceable Big-Digit Type K Thermometer). A *wild-type* control was run at the beginning of every data collection session. For the experiment, a fly of known sex was placed under the plastic cover and a Styrofoam box was placed to cover the experimental area to remove light and allow for the experiment to be conducted in dim ambient light (<10 lux). The flies were given 2 minutes to acclimate on the 25± 1°C side. A HERO8 GoPro was positioned at the top of the Styrofoam box (8 inches in height). The motion of the fly was recorded by taking a time-lapse photo every second for 120 seconds.

Adult optogenetic assays were recorded by a HERO8 GoPro. Half of a clear plastic cover was covered with black tape. A piece of white paper was put under the cover to reduce noise from the background. A Styrofoam box was used to cover the experimental area. A HERO8 GoPro was mounted at the top of the Styrofoam box. On one side of the GoPro, a red-light source (Tyrrell et al., 2021) was attached. The plastic cover was placed on the other side of the GoPro to avoid the glare that is caused by the direct light. A single fly of known sex was pipetted under the plastic cover and given 2 minutes to acclimate. The light source was turned on and the motion of the fly was recorded by taking a time-lapse photo every second for 60 seconds. Positive and negative controls were run at the beginning of every data collection session.

### Larval behavioral assays

Flies were maintained at 25°C under 12-hour light/12-hour dark cycles. Larvae were collected as described with some modifications (Tyrrell et al., 2021). Briefly, each vial contained 20-45 males and females. These flies were given at least 24 hours to recover from CO_2_ before being tapped over to new vials containing yeast granules. They were allowed 4 to 8 hours to lay eggs and, on day 4, larvae were collected using 10 mL of 20% w/v sucrose solution. Larvae were collected after 20 minutes and thoroughly washed three times with diH_2_O. Larvae were then plated on a 60 mm tissue culture dish (Corning) with about 13 mL of 3% room temperature (about 20°C) agar gel and given 5 to 10 minutes to recover from the washing process and acclimate to the agar.

The two-choice assay was performed as described with some modifications (Tyrrell et al., 2021). Two steel plates on different hot plates were separated by 1/16 inches (the release zone). For the free movement behavioral assay, a steel plate was placed on a hot plate and its surface temperature was adjusted to 18± 1°C. A matte black poster sheet was placed on top of steel plates. A plastic sheet protector was placed on the poster sheet to prevent warping from moisture. A 3% agar gel (10 × 9.5 inched) was placed on the plastic protector sheet and evenly positioned at the release zone. The surface temperature was 18± 1°C on one side of the gel and 25± 1°C on the other. The release zone was labeled at the top and bottom of the gel. The temperature was monitored before each trial using a surface temperature probe (80PK-3A, Fluke) and thermometer (Fisherbrand Traceable Big-Digit Type K Thermometer). A *wild-type* control was run at the beginning of daily experiments. Water was gently sprayed between trials to moisten the agar surface. A larva was placed at the release zone and given 2 minutes to wander. The experiment was conducted at dim ambient light (<10 lux). A GoPro was suspended above the gel (10.5 in) to record the motion of the larva for each trial.

Larval optogenetic assays were recorded by a Sony HDR-CX405 camcorder with the internal infrared filter removed and an 830 nm long-pass filter (FSQ-RG830, Newport) installed. A 3% agar gel was cut to 3 × 5 inches. A sheet of a matte black poster was placed under the gel to reduce background noise signals and infrared light (4331910725, Amazon) was used to visualize the larvae. Two red-light sources (Tyrrell et al., 2021) were attached 10 inches above the gel at a 45° angle on both sides of the gel such that the light intensity was even throughout the gel (∼3 klux) and no glare was created. A third red-light source was placed in a box outside the experimental setup but within the view of the camera to serve as an indicator to record when red lights were on or off. Larvae were collected and prepared for the assay as detailed above except that they were kept in food with 40 μM all *trans*-retinal (ATR, Sigma-Aldrich) and in dark for 72 hours. An individual larva was placed on the agar gel and given 30 seconds to acclimate. For the recording, the larva was given 30 seconds to wander followed by 3 cycles of 5 seconds shone under red lights and a 15 second recovery period.

### Preprocessing photos for adult behavioral assays

The individual photos for each trial were combined into a single file using Fiji and converted to 8-bit grayscale (File > Import > Image Sequence function; boxes of Convert to 8-bit Grayscale and Sort names numerically were selected) (Schindelin et al., 2012). For two-choice and optogenetic assays, the Rotate feature (Image > Transform > Rotate) was used to rotate images until the steel plate dividing line or tape line was shown as completely vertical. A black line was drawn along the steel plate boundary to separate the two steel plates (temperatures) using the draw function (Edit > Draw) and applied to all images. Then, for all the assays, the Crop feature (Image > Crop) was used to only keep the experimental area. Next, backgrounds were subtracted from all images (Process > Subtract Background; set rolling ball radius of 20.0 pixels; the box of light background was selected). The brightness/contrast was adjusted to enhance the difference between the dark fly and the white background (Image > Adjust > Brightness/Contrast). Lastly, the threshold was set to the automatic suggested setting (Image > Adjust > Threshold; Default and B&W settings were chosen, the box of Don’t reset range was selected). In the Convert Stack to Binary box, the method was set to default, background was set to light, and the box corresponding to calculate threshold for each image was selected. The preprocessed image was then saved as a TIFF (File > Save as > TIFF) for analysis.

### Preprocessing videos for larval behavioral assays

Videos were converted to .avi and the resolution was decreased to 760×480 by Any Video Converter 9 (AnvSoft). They were then uncompressed by the command line tool ffmpeg to be compatible with Fiji.

Videos were imported to Fiji and converted to 8-bit grayscale (File > Import > AVI; the box of Convert to Grayscale was selected). For two-choice assays, the Rotate feature (Image > Transform > Rotate) was used to rotate images such that the marker line indicating release zone was completely vertical. Along the release zone, white lines were drawn at the top and bottom of the larval motion zone; they must be close to but not in the larval motion zone and applied to the first image only (Edit > Draw). Then, an area that was slightly larger than the larval motion zone and included the top and bottom white lines was selected and the Crop feature (Image > Crop) was applied for all images. Next, background was subtracted from all images (Process > Subtract Background; rolling ball radius of 50.0 pixels). Finally, the brightness/contrast was adjusted to enhance the difference between the white larva and the black background (Image > Adjust > Brightness/Contrast; set Maximum to the left and applied; then set Contrast to the right and applied). The preprocessed image was saved as a .avi file (File > Save as > AVI) for analysis.

### TrackMate Analysis

TrackMate in Fiji was used to analyze the moving distances and preference/avoidance indices (Tinevez et al., 2017). Preprocessed .tif files or .avi files were opened by Fiji and TrackMate was run. In the box of the LoG Detector, we suggested to adjust parameters of Estimated blob diameter, Threshold, and Median filter. For adult flies, set Estimated blob diameter to 27.0 – 40.0 pixels, Threshold to 1.0 - 2.5, and select Median filter. For larvae, set Estimated blob diameter to about 10.0 pixels, Threshold to 1.0, and deselect Median filter. In the box of Set filters on spots, filters could be used to remove aberrant ROIs by choosing the X and Y region for ROIs as well as the Quality of the ROIs found. Alternatively, aberrant ROIs can be removed manually from the resulting .csv file. In the box of the Simple LAP tracker, the Linking max distance, the Gap-closing distance, and the Gap-closing max frame gap were suggested to adjust. For adult flies, set the Linking max distance, the Gap-closing distance to 1500 pixels and the Gap-closing max frame gap to 2. For larvae, set the Linking max distance, the Gap-closing distance to 25 pixels and the Gap-closing max frame gap to 2. In the box of Select an action, Export all spot statistics was selected and Execute was clicked. The All Spots statistics file was cross-referenced with the spot detection in Fiji to ensure only one spot was marked for each time stamp, with erroneous duplicates and/or aberrant ROIs deleted. Then the All Spots statistics was saved as .csv files.

The moving distance from frame n to the next frame was calculated through the following formula:

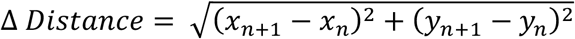

The Preference Index (PI) or Avoidance Index (AI) was calculated by using the X position. The PI for the two-choice assay was calculated based on the time the animal spent in each temperature zone and using the following formulas:

For adults:

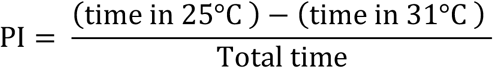

For larvae:

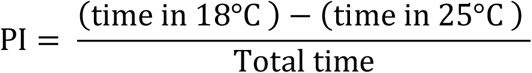

The AI for the adult optogenetic assay was calculated based on the time a fly spent in dark or red light and using the following formulas:

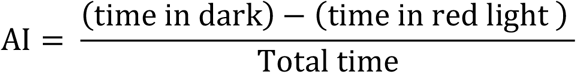

### Statistical analysis

Statistical details of experiments are mentioned in the figure legends. The normality of distributions was assessed by the Shapiro-Wilk W test (*p* ≤ 0.05 rejected normal distribution). Statistical comparisons of normally distributed data were performed by the Welch’s t test. For data that did not conform to a normal distribution, statistical comparisons were performed by the Mann-Whitney test. Data analysis was performed using GraphPad Prism 9.

## Data availability

Original statistics and raw data are available at: https://doi.org/10.7910/DVN/SNBQC2.

---

**Movie 1. Representative trajectories of *Drosophila* adult and larval behaviors**. The following behaviors are included: the *wt* and *Gr28b*^*MB*^ single-fly free movement and two-choice thermotactic assays, *HC*>*CsChrimson* single-fly optogenetic assay (without and with dietary ATR), and *wt* and *Ir93a*^*MI*^ single-larva free movement and two-choice thermotactic assays.

